# Kinetic frustration by limited bond availability controls the LAT protein condensation phase transition on membranes

**DOI:** 10.1101/2021.12.05.471009

**Authors:** Simou Sun, Trevor GrandPre, David T. Limmer, Jay T. Groves

## Abstract

LAT is a membrane-linked scaffold protein that undergoes a phase transition to form a two-dimensional protein condensate on the membrane during T cell activation. Governed by tyrosine phosphorylation, LAT recruits various proteins that ultimately enable condensation through a percolation network of discrete and selective protein-protein interactions. Here we describe detailed kinetic measurements of the phase transition, along with coarse-grained model simulations, that reveal LAT condensation is kinetically frustrated by the availability of bonds to form the network. Unlike typical miscibility transitions in which compact domains may coexist at equilibrium, the LAT condensates are dynamically arrested in extended states, kinetically trapped out of equilibrium. Modeling identifies the structural basis for this kinetic arrest as the formation of spindle arrangements, favored by limited multivalent binding interactions along the flexible, intrinsically disordered LAT protein. These results reveal how local factors controlling the kinetics of LAT condensation enable formation of different, stable condensates, which may ultimately coexist within the cell.

## Introduction

Protein condensates are emerging as one of the key organizational features in cell biology^[1,2]^. Widely studied liquid condensates of proteins and nucleic acids are often described in terms of liquid-liquid phase separation (LLPS)^[1,3,4]^. Recently, a distinct class of protein condensates, consisting of precisely regulated assemblies of signal transduction proteins on the membrane surface, have been identified^[5-7]^. These two-dimensional condensates form through a multivalent bond percolation network of specific protein-protein interactions, a common motif of which is the modular binding interaction between SH2 domains and phosphorylated tyrosine residues. The crosslinking protein-protein interactions are relatively strong (a few tens of *k*_*B*_T) and occur through a limited number of specific binding sites. This limited bond availability establishes a distinct physical difference with the three-dimensional protein-nucleic acid condensates, which are mediated through large numbers of weak interactions^[1-3]^. Under physiological conditions in the living cell, selective kinase and phosphatase reactions regulate phosphorylation at the tyrosine binding sites and, in this way, govern the formation and dissolution of the two-dimensional signaling protein condensates. Controlled tyrosine phosphorylation on scaffold and signaling proteins is a central feature of information processing within cellular signal transduction networks, positioning these membrane-associated condensates under direct control of signal transduction pathways^[5-7]^.

Here, we describe studies of phase transition kinetics in a system consisting of Linker for Activation of T cells (LAT), Growth Factor Receptor-Bound Protein 2 (Grb2), and Son of Sevenless (SOS). LAT is an intrinsically disordered, single-pass membrane anchored protein that—together with Grb2, SOS and other molecules—serves as a hub for signal processing in the T cell receptor (TCR) signaling pathway (Figure 1a)^[8,9]^. LAT is a substrate for selective phosphorylation by the Zap70 kinase in response to TCR activation by antigen. LAT has nine tyrosine sites, at least 3 of which can readily bind Grb2 through its SH2 domain when phosphorylated. Grb2 additionally has two SH3 domains, which can bind the proline-rich (PR) domain on SOS. SOS is capable of binding more than one Grb2 in this way, and thus provides a crosslinking mechanism that leads to networked assembly of a LAT:Grb2:SOS protein condensate on the membrane surface^[5,7]^. Other crosslinking interactions, such as a direct Grb2:Grb2 dimer interface^[10]^, may participate but Grb2-mediated crosslinking is primarily limited to three LAT tyrosine sites—thus only achieving the minimum valency required for bond percolation. Another tyrosine site on LAT, which exhibits differential phosphorylation kinetics^[11]^ and selectivity for the key signaling molecule PLCγ[12], provides a fourth crosslinking pathway that may be physiologically important in nucleation of the LAT condensate^[13,14]^. Individually, the LAT:Grb2 and Grb2:SOS interactions exhibit relatively fast binding kinetics^[15]^. However, rapid rebinding in the condensed state substantially extends the dwell time distribution for individual molecules, and has been identified as a functional mechanism by which the condensate can control signaling output^[7]^. The discrete binding interactions to specific sites, complex molecular kinetics, and intrinsic two-dimensionality suggest that LAT condensates are likely to exhibit very different material properties compared with three-dimensional protein-nucleic acid condensates^[1-4,16]^.

**Figure 1.**
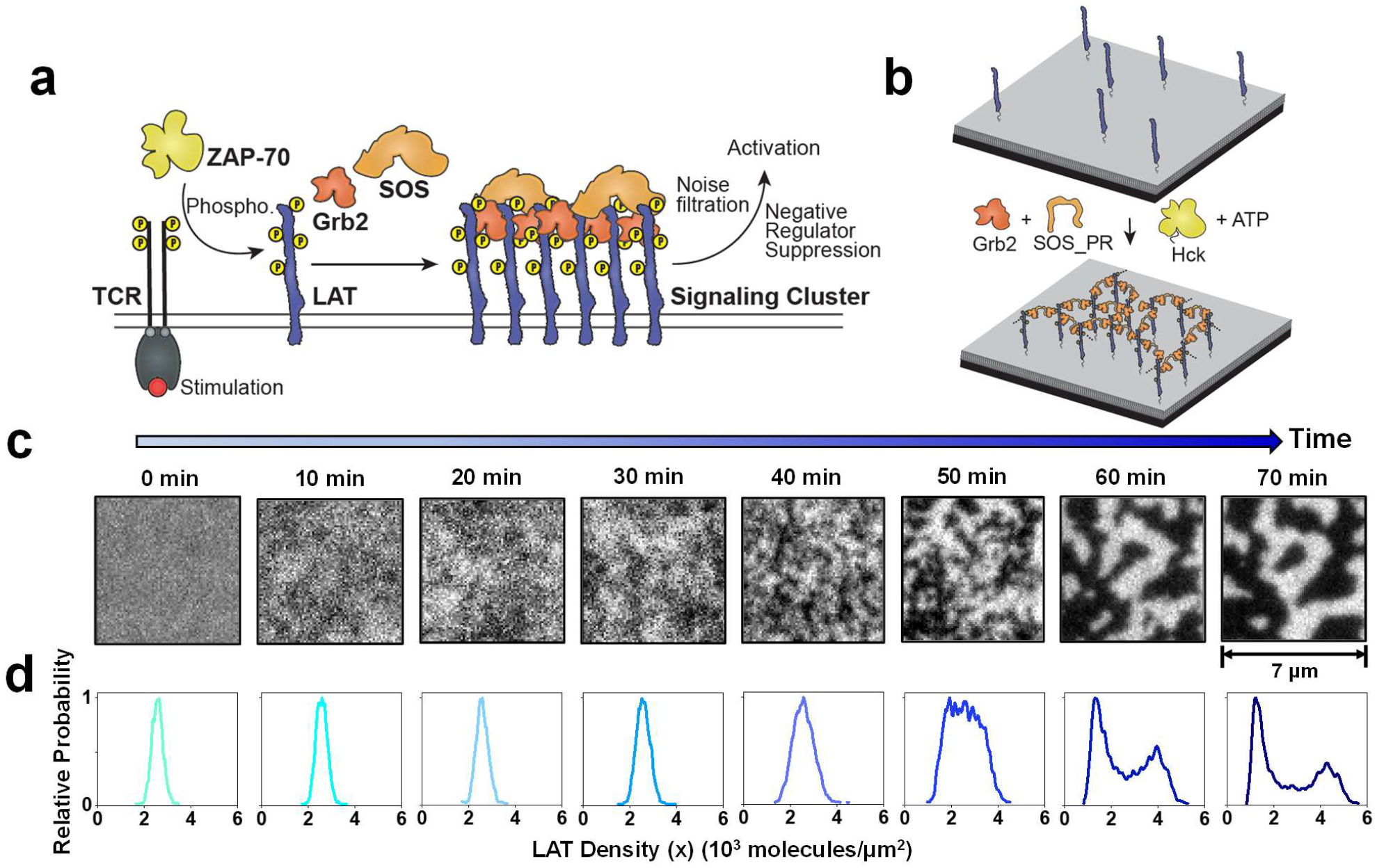
Schematic illustration of the LAT:Grb2:SOS_PR condensation phase transition system. a) The TCR signaling pathway which involves assembly formation with LAT:Grb2:SOS. b) In vitro reconstitution of the LAT:Grb2:SOS_PR condensation phase transition on a supported lipid bilayer, where the assembly eventually relaxes to a disordered gel state. c) TIRF images of the LAT-555 molecules during the progress of a phase transition. d) Corresponding LAT density (x) distribution over time. The density of LAT molecules was calculated based on TIRF intensity and FCS readout calibration.

We reconstituted the LAT:Grb2:SOS system from purified components and performed detailed microscopic imaging studies tracking kinetics of the phase transition process on supported membranes. The results reveal that LAT:Grb2:SOS condensates undergo slow coarsening dynamics and kinetic arrest, preventing the condensate from reaching equilibrium. Temperature dependence of the phase transition kinetics, changes in molecular mobility, and the apparent final state composition exhibit signatures of a kinetically dominated control mechanism. We introduce a coarse-grained model of the LAT condensation based on the modular molecular binding kinetics, valency, and two-dimensional mobility constraints that proved capable of recovering key experimental observations. The modeling reveals that limited valency and intrinsic flexibility of LAT favors the formation of spindle-like structures that act as kinetic traps, substantially slowing further equilibration of the system and leading to dynamic heterogeneity. We further discuss how insights from these experiments and modeling studies prompt some alternative interpretations of the mechanistic role of LAT and its condensation phase transition in T cell signaling.

## Results

The LAT protein condensation phase transition, which naturally occurs downstream of T cell receptor signaling during T cell activation, can be reconstituted on supported lipid bilayers (SLBs) from purified proteins (Figure 1a and 1b)^[5,15,17]^. Briefly, AlexaFluor-555 labeled His6-LAT (residues 27-233, LAT-555) was anchored on the SLB through Ni^2+^-histidine chelation with Ni^2+^-NTA-DOGS lipid, doped into the membrane. Prior to condensation, LAT molecules were homogeneously distributed and diffuse freely as monomers (D = 1.6 ± 0.3 μm^2^/s) on the membrane surface, as observed with total internal reflection fluorescence (TIRF) microscopy imaging and fluorescence correlation spectroscopy (FCS) (Figure S1). In these studies, LAT is maintained in its phosphorylated state by the Src family kinase Hck, which is also tethered to the membrane and can phosphorylate LAT to completion^[15]^. To initiate the condensation phase transition, phosphorylated LAT on the membrane was first allowed to incubate with Grb2 for 20 minutes, which enables binding of Grb2 to phosphorylated tyrosine sites on LAT to come to equilibrium. Under the concentration conditions used in these experiments, Grb2 binding alone is insufficient to induce condensation and the LAT remains homogeneous (Figure S2). To trigger the phase transition, the proline-rich (PR) domain of SOS (here simply referred to as SOS) was introduced to the Grb2 in solution. The SOS PR domain strongly crosslinks Grb2 and this is sufficient to trigger the LAT:Grb2:SOS condensation^[5,7,15]^. After abrupt introduction of SOS, the ensuing LAT condensation phase transition is tracked through time using TIRF imaging (Figure 1c).

The LAT:Grb2:SOS system undergoes a macroscopic phase transition to reach a steady state with two types of domains of drastically different LAT densities. Regions with high (bright) and low (dark) LAT concentration, defined as condensed and dispersed phases, respectively, are clearly visible. The condensed domains are quasi-stationary with stable and non-circular geometry, as well as with a broad range of sizes, and they exhibit constant local fluctuations at the edges (Movie S1). No active (ATP consuming) process is required to maintain the size and morphology of the condensed domains. In these experiments, the phase transitions occur over timescales of ∼1 hour, but this timescale is related to the macroscopic size of the system. Over shorter length scales (e.g. a few hundred nanometers for some LAT condensates imaged in cells), timescales will be significantly faster. We quantify the state of condensation by measuring the LAT surface density distribution on the membrane from the calibrated pixel brightness histogram of the fluorescence images (Figure 1d, analysis details are provided in the SI and Figure S3). Each image pixel samples an area of ∼ 9400 nm^2^ on the membrane. Before condensation, the LAT density distribution is well fit by a single Gaussian peak with a narrow variance consistent with a random distribution of monomeric LAT. Over the course of the phase transition, the distribution broadens notably after an initial lag time, and ultimately converges into two prominent and well separated peaks (Figure 1d).

We first characterize the final state under a variety of conditions to map the apparent LAT condensation phase diagram. To facilitate this analysis, the steady-state LAT density distributions were fit to two Gaussian peaks, the positions of which correspond to the average LAT density in the condensed and the dispersed phases (*c*_c_ and *c*_d_, Figure 2a). This fitting procedure was used to map the LAT density difference in the two phases across a variety of experimental temperatures to establish a phase diagram (detailed description of the analysis provided in the SI and Figure S4). For purposes of comparison, the phase coexistence boundary of a classical miscibility phase diagram is sketched in Figure 2b. For system compositions at temperatures below the phase coexistence line, condensed and dispersed phases coexist with compositions marked by the dashed tie lines. For equilibrium miscibility phase separation, these coexisting phase compositions correspond to thermodynamic free energy minima and are fixed; varying initial protein concentrations leads to differing amounts of the two phases but their compositions are preserved^[18]^. The coexistence curve (or binodal) is indicated on Figure 2b by the solid black line, identifying specific coexisting phase compositions at each temperature (highlighted with the horizontal dashed tie-lines, for examples). Generally, at higher temperatures the coexisting phases become more similar in composition until the critical temperatures (*T*_c_) is reached, above which the system remains macroscopically homogeneous. The experimentally measured phase coexistence diagram for the LAT:Grb2:SOS condensation phase transition is shown in Figure 2c. For temperatures closer to the physiological temperature (≥ 31 °C), the coexisting phase compositions resemble a classic miscibility phase diagram. However, at lower temperatures, the experimental data deviate notably, with coexisting phases apparently much closer in composition. Furthermore, the two-dimensional phase transition does not follow the classical tie-line principle with fixed *c*_c_ and *c*_d_. Instead, LAT concentrations in the two phases grew correspondingly with increasing initial LAT concentrations (Figure 2d), and the ratio of *c*_c_ to *c*_d_, known as the partition coefficient (*K*), remained constant (Figure S5). These results indicate that the apparent final state of the LAT:Grb2:SOS phase transition is not an equilibrium free energy minimum, instead, it is kinetically and stably trapped in an out-of-equilibrium state.

**Figure 2.**
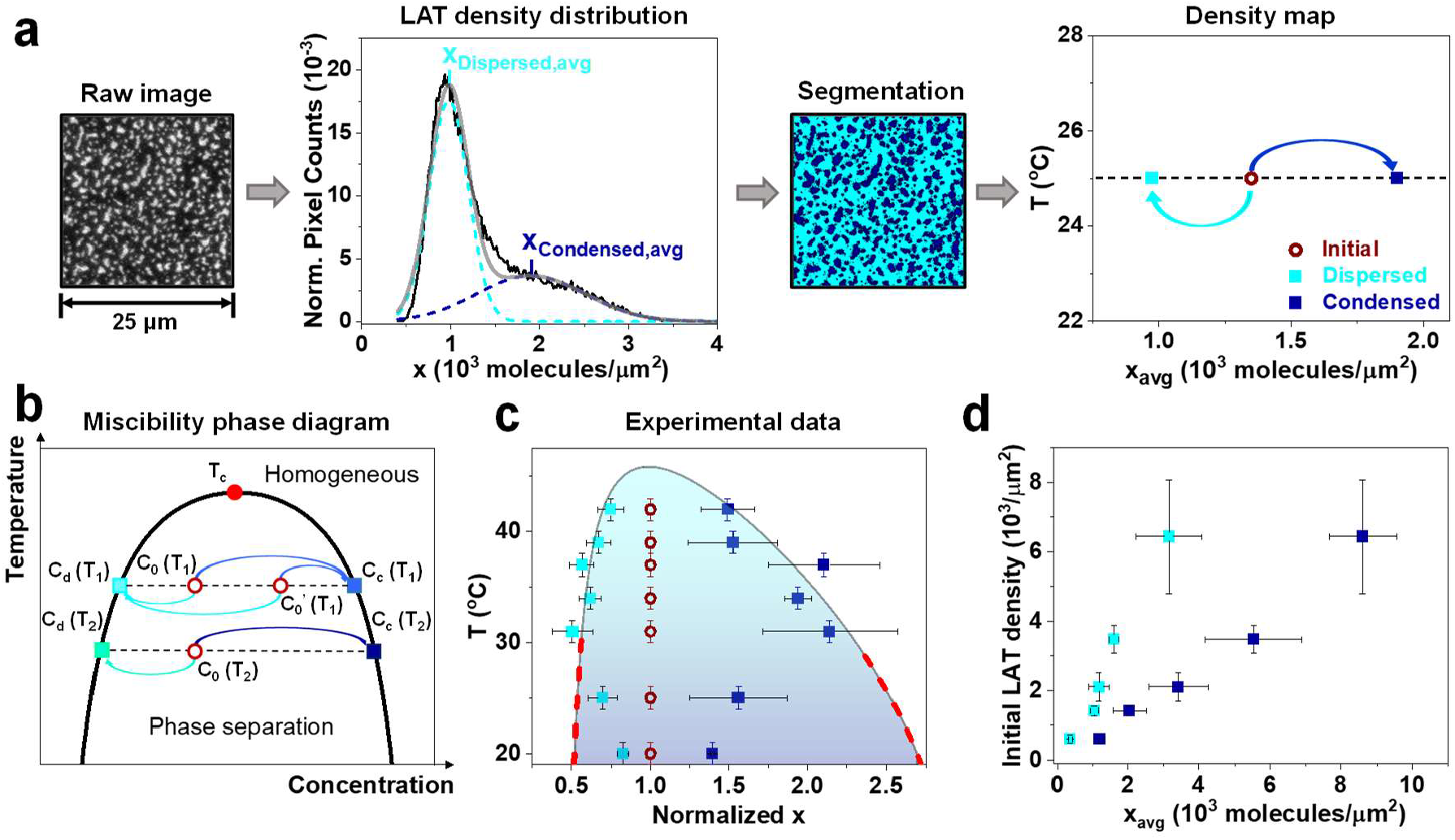
Density map of the LAT:Grb2:SOS_PR system after the phase transition reaches the apparent final state. a) Illustration of the analysis strategy to obtain the density map. The calibrated LAT density distribution from the raw image was fitted with two Gaussian peaks, the positions of which correspond to the average LAT densities in the dispersed and the condensed phases. A representative density map at 25 °C is shown in the right panel. Red open circle: initial LAT density before phase transition, royal blue solid square: average LAT density in the condensed phase, cyan solid square: average LAT density in the dispersed phase. b) Demonstration of a classical miscibility phase diagram, where the condensed phase and the dispersed phase are at equilibrium. c) Normalized average LAT density in the condensed domains and in the dispersed domains as a function of experimental temperature at an initial LAT density of 1200 ± 600 molecules/μm^2^. The shaded area corresponds to the two-phase region of a modeled miscibility phase diagram. The red dashed lines on the binodal curve indicate deviation from the experimental data. d) Average LAT density in the condensed and the dispersed phases plotted on the x-axis with different initial LAT density plotted on the y-axis.

We analyzed the kinetics of condensation by tracking the LAT surface density distribution over the course of the phase transition. The normalized variance of this distribution, corresponding to the mean square LAT density fluctuation (with spatial sampling set by the ∼ 9400 nm^2^ image pixel size), provides a simple and monotonically increasing measure of the extent of phase separation. Kinetic traces of this parameter measured for LAT:Grb2:SOS phase transitions at a variety of temperatures are plotted in Figure 3a. The progress curves qualitatively exhibit a lag stage and a growth stage, in which small and dynamic density fluctuations precede a macroscopic structural reorganization.

**Figure 3.**
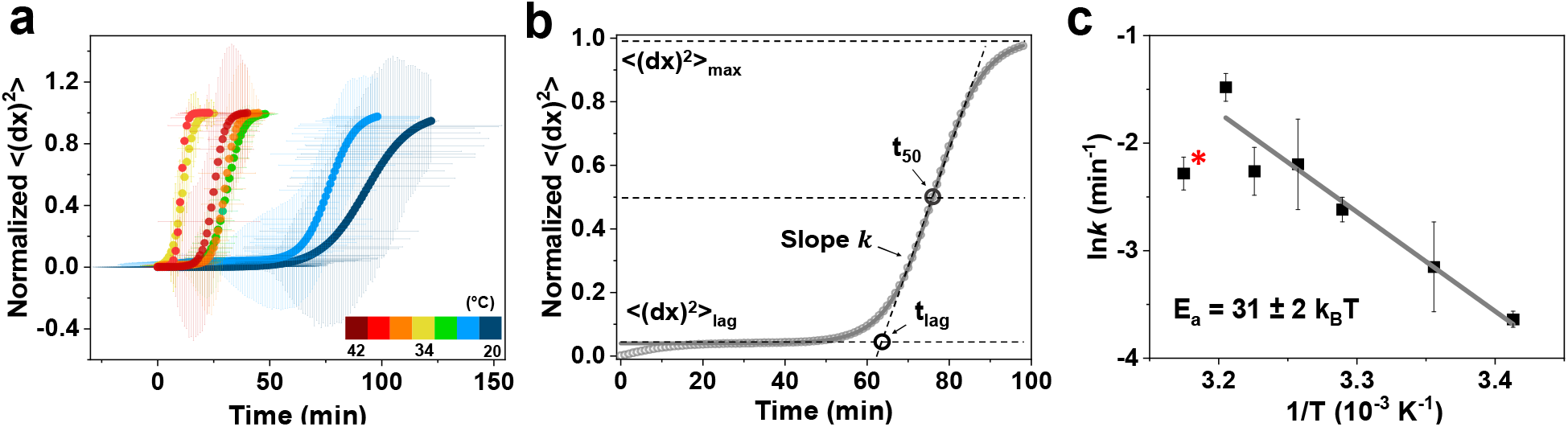
Kinetic characterizations of the LAT:Grb2:SOS_PR condensation phase transition. a) Normalized mean square LAT density fluctuation ((dx)^2^>) during the phase transition in a temperature range from 20 °C to 42 °C. The data are color-coded according to the temperatures. Error bars represent SD from at least three repeats of experiments at each temperature. b) Illustration of the analysis strategy to extract the phase transition rate (k) and lag time (t_lag_)by fitting the kinetic profile to a sigmoidal function. c) k in log-scale as a function of 1/T fitted to the Arrhenius equation (20 °C to 39 °C, the fit did not include the 42 °C data marked with asterisk), with a calculated E_a_ = 31 k_B_T ± 2 k_B_T.

An empirical sigmoid model has been previously used to characterize similar behavior in the dynamics of kinetically controlled phase transitions of amyloid formation^[19,20]^. From such analysis of the LAT transition kinetics, the principal parameters for the transition rate (*k*) and the lag time (*t*_lag_) were extracted (Figure 3b), where *k* characterizes the rate of the macroscopic structural rearrangement at *t*_50_ and *t*_lag_ represents the time delay between adding linker proteins to the start of the macroscopic phase transition (see SI for further discussion). Increasing temperature from 20 °C to 39 °C progressively reduced an apparent kinetic bottleneck and sped up the phase transition, as quantified by increasing values of *k* and decreasing values of *t* _lag_ in the sigmoid model fits. This temperature dependence is counter to expectations from supersaturation-driven miscibility phase separation kinetics, where increasing temperature is expected to decrease the thermodynamic driving force for condensation and thus slow the rate of transition^[21,22]^. Instead, the observed kinetic feature is reminiscent of the non-equilibrium phase transitions, such as gelation and glass transitions^[21,23]^.

Over the 20 °C to 39 °C temperature range, the LAT:Grb2:SOS phase transition exhibits Arrhenius kinetics with an activation energy *E*_a_ ∼ 31 *k*_B_*T* (Figure 3c). This energy scale closely matches the estimated bonding energy between LAT molecules (see SI for details), which suggests that the phase transition is rate limited by bond formation and breakage. An additional feature in these results is that the phase transitions kinetics exhibit a non-monotonic dependence on temperature, with the fastest rate occurring at an intermediate temperature of 39 °C. At the highest temperature measured (42 °C, marked with asterisk in Fig. 3c), the kinetic barrier has largely been eliminated but an attenuated thermodynamic driving force for phase separation could explain the observed decreased *k* and overall slower transition kinetics. It is worth noting that, over the same range of temperature, there is negligible change in the unassembled LAT mobility (Figure S6); the diffusive mobility of LAT itself is likely not a significant contributor to the phase transition kinetic behavior.

To investigate the microscopic structures within the protein network during the phase transition and maturation process, we conducted single molecule tracking experiments on LAT. Briefly, 0.1 mol% AlexaFluor-488 labeled LAT (LAT-488) was doped into 99.9 mol% LAT-555, allowing visualization of both the macroscopic LAT configuration as well as individual molecular mobilities within. Four different stages of the macroscopic phase transition process, as depicted in Figure 4a, were characterized: (*i*) before adding Grb2 and SOS_PR; (*ii*) the lag stage; (*iii*) the growth stage; (*iv*) the apparent final state. Prior to any condensation, LAT exhibits simple Brownian motion with diffusion coefficients of ∼1.1 μm^2^/s (Figure 4b). At later stages during the transition, a significant amount of LAT exhibits highly trapped motion and some LAT exhibits intermediate frustrated motion (Figure 4c).

**Figure 4.**
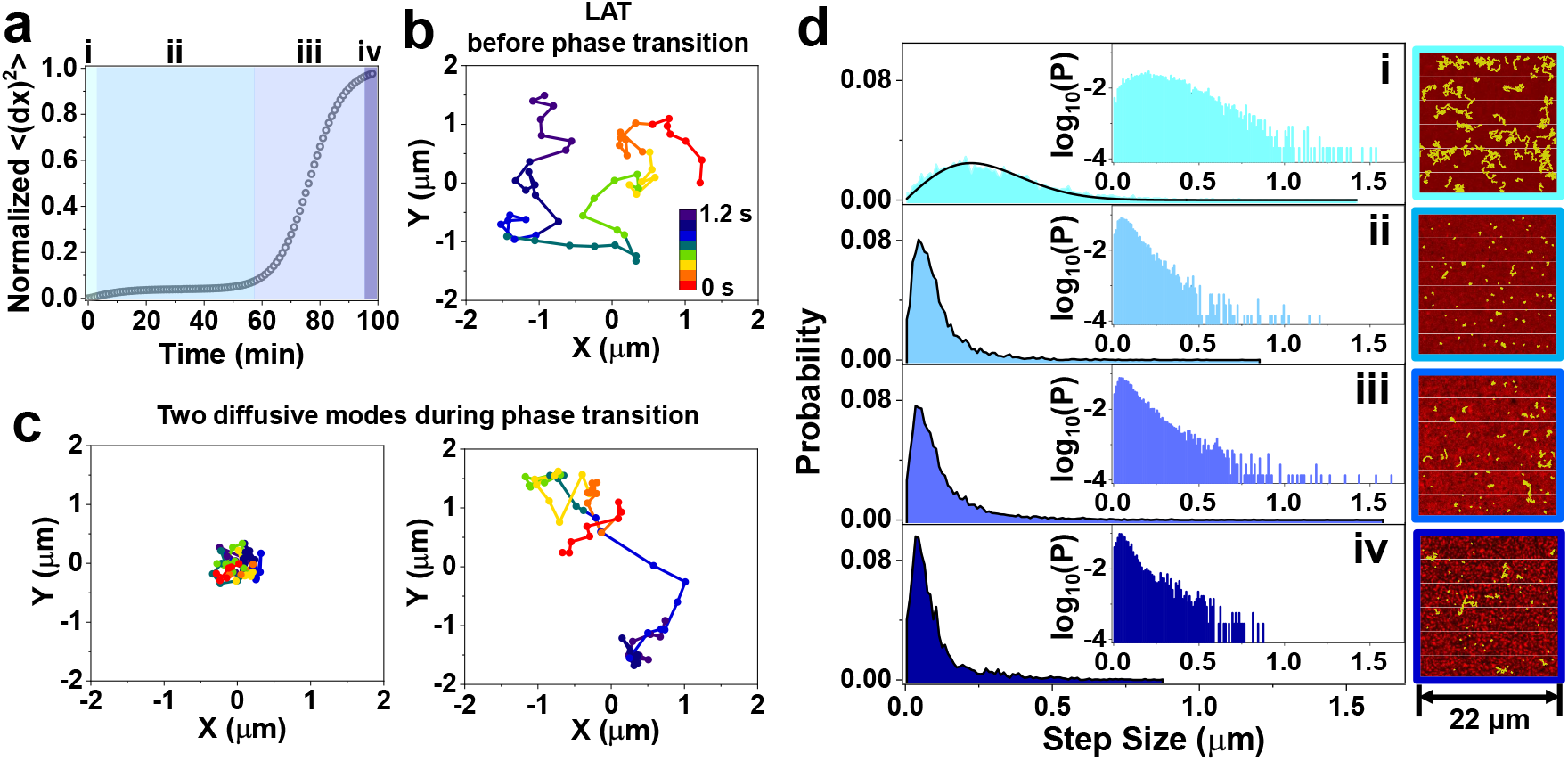
Single-molecular dynamics during the LAT:Grb2:SOS_PR condensation phase transition. a) Representative kinetic profile of the phase transition recorded on the 561 nm excitation channel (99.9 mol% LAT-555), with four different stages of the phase transition color coded (i. before phase transition, ii. lag stage, iii. growth stage, iv. apparent final state) for the single molecule tracking experiments on the 488 nm excitation channel (0.1 mol% LAT-488). b) & c) Representative tracking trajectories of LAT-488 before phase transition and during phase transition, where LAT displayed at least two drastically different mobility. The time axis is color-coded in the trajectories. d) Step size distribution of the mobile LAT-488 molecules at the four stages and corresponding representative phase transition configurations (red color represents LAT-555) and tracking trajectories (yellow color represents LAT-488). Inset: same step size distribution with probability plotted in log scale. The data in stage i was fitted to a single-component Rayleigh distribution. 400 ± 100 molecules were tracked at 33 Hz for 100 frames to generate each distribution profile.

In Figure 4d, plots of measured LAT step size distributions during the four stages are displayed, along with the corresponding macroscopic phase transition images and single-molecule tracking trajectories. In stage i, the homogeneously distributed and unassembled LAT molecules displayed simple Brownian motion and are well fit by a Rayleigh distribution for single component diffusion. In stages ii, *iii*, and *iv* the data shifts to smaller step sizes with the majority the molecules being highly constrained and localized. The molecular motion is highly non-Brownian and a significant amount of heterogeneity exists. The largely reduced LAT mobility in the lag stage suggests that the LAT:Grb2:SOS bonding network readily formed oligomeric complexes early on. These structures are expected to contribute to the small LAT density fluctuations observed at the ensemble level, and they are kinetically stable possibly due to local valency saturation^[24]^. In stage *iii*, surprisingly, the LAT step size distribution broadened slightly. Despite the majority of LAT molecules still exhibiting trapped motion, we observed an increased population of fast LAT with large step size compared to stage *ii*. This minor fast-moving population was observed to jump between two condensed domains, and it is potentially the major dynamic driving force for the macroscopic structural rearrangement during the slow coarsening process. In stage *i*v, the step size distribution narrowed down again. The number of LAT molecules in trapped states within condensates further increased. Interestingly, the LAT molecules observed in the dispersed phase within the tracking timescale displayed smaller step size than the freely diffusing LAT in stage *i*, possibly indicative a stably formed dimers or small oligomers. Note that two LAT molecules that are triply bound to each other will not readily form bonds with other LAT molecules due to limited valency, and would be relatively stable as a free dimer.

In order to understand the slow coarsening kinetics accompanying LAT aggregation, we have developed a coarse-grained model that encodes only general biophysical processes. We model membrane-bound monomer LAT molecules as diffusing in two-dimensions with thermally driven Brownian motion. Each molecule excludes a volume with a characteristic diameter, **σ**, determined by the linear polymer’s radius of gyration, which is estimated to be nearly **σ=5** nm and is implemented with a repulsive pair potential. Most of our studies are considered at a density of 400 molecules/μm^2^. We envision the bonding of a Grb2:SOS_PR:Grb2 complex to two adjacent LAT molecules as an effective first order process, which is valid in the limit of a well-mixed solution phase and fast binding kinetics of SOS_PR:Grb2. The Grb2:SOS_PR:Grb2 linker is bound with a attempt rate *k*_b_, with unbinding kinetics obeys detailed balance with respect to a bonding potential that includes a minimum bond energy, −*Ē*_b_, and is held at a rest length of 15 nm with a stiff harmonic potential. Additionally, we impose a kinetic constraint that each LAT can form at most three bonds with other LAT molecules. The action of the linker is to bond LAT molecules together, co-localizing them in space. A phase diagram analogous to Figure 2c is shown in Figure S7, in which at elevated temperatures LAT is found to be uniformly dispersed, while at low temperatures LAT molecules aggregate into a condensed gel that lacks long range positional order consistent with experiments. The phase separated temperature is set by the bond energy, and the disorder results from the formation of a network with sufficiently few mechanical constraints that there are many low energy deformation modes.

The combination of diffusion and complex formation with limited LAT valency is sufficient to reproduce many of the experimental observations in Figure 2-4. Specifically, we find an initially dispersed system when quenched below the phase separation transition temperature grows through a coarsening process in which an initial increase in LAT bonding occurs quickly and only over much longer timescales do isolated clusters aggregate. The initial bonding occurs with a characteristic time identified with the single complex formation rate, while the growth of large compact clusters occurs two orders of magnitude slower (Figure 5a). Clusters grow as a power-law in time with a growth exponent comparable with the experimental observations (Figure 5a and Figure S8). At short times, the increase in the number of bonds occurs through the growth of many small clusters (Figure 5d). The limited valency results in the formation of extended spindle-like structures at intermediate times. These spindle-like structures are traps with long structural reorganization times due to limited bond availability within them. The spindle-like structures condense over time and display a meta-stable microphase that is kinetically trapped where multiple aggregates exist. At the longest times, the system eventually forms a single large cluster, mediated by rare bond breaking events. The resulting gel exhibits large density fluctuations and a mean density far from close packed. The properties of the transient multi-droplet system depend on the ratio of the rate of bond formation, and the diffusion of LAT molecules (see more detailed discussion about the steady state properties in Figure S9).

**Figure 5.**
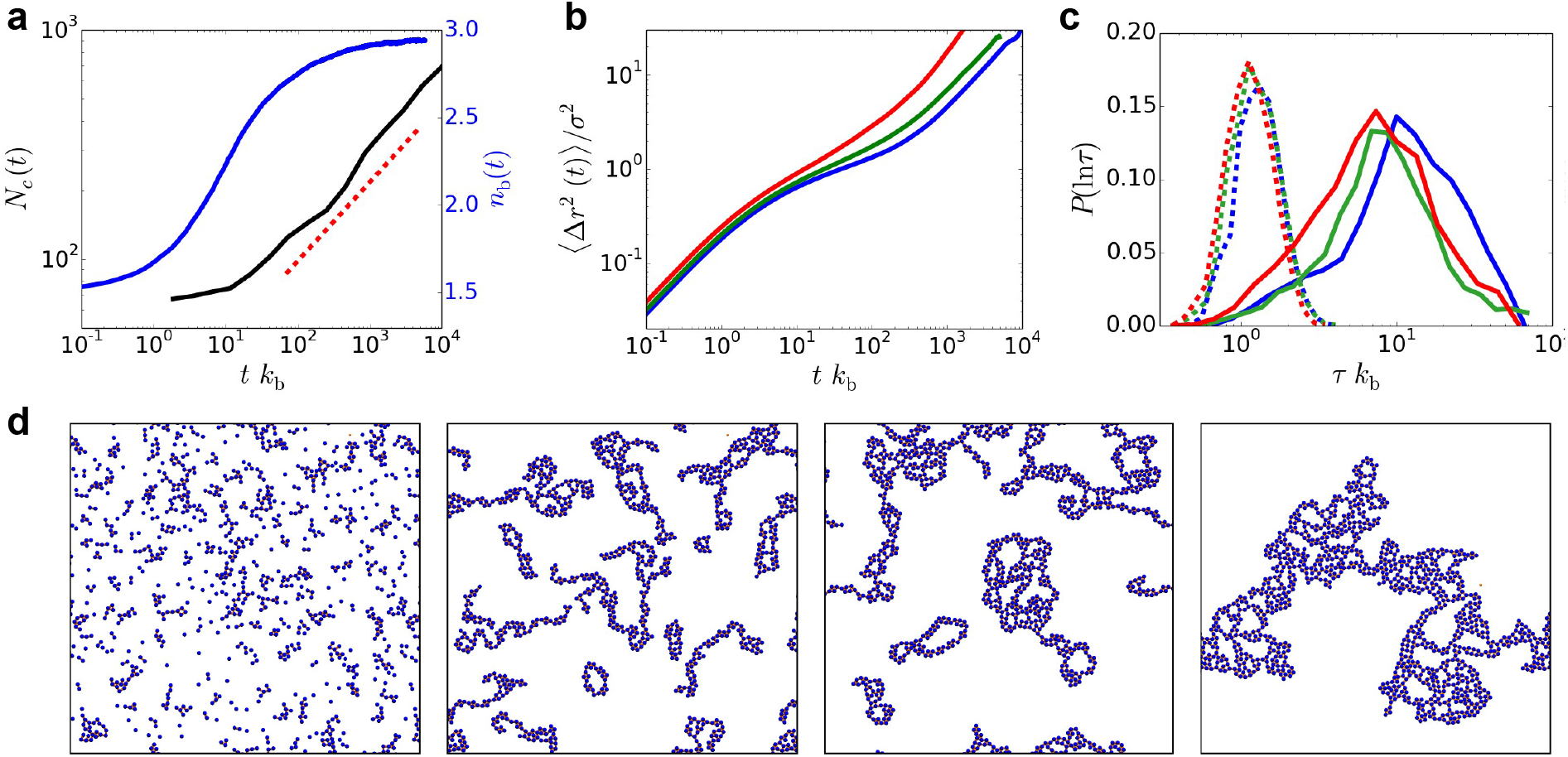
Simulation results of LAT model. a) The largest cluster size (black line) and average bonds (blue line) as a function of time, with comparison to a 1/3 coarsening power law (red dashed line) for a reduced density of ρσ^2^ = 0.1 and a reduced temperature of k_B_T/Ē _b_ = 0.15. b) The mean squared displacement of the dense phase as a function of time showing caging effects that become more distinct with decreasing temperature for reduced temperatures of k_B_T/Ē _b_ = 0.18 (red), 0.15 (green), and 0.14 (blue) and reduced densities ρσ^2^ = 0.27, 0.32, and 0.36. c) The distributions of persistence times (solid line), the time to move their particle diameter, σ, and exchange times (dashed line), the time to move another distance σ given that a LAT molecule has already moved that distance, for the same three temperatures and densities in b). The lack of overlap demonstrates dynamic heterogeneity. d) Typical trajectory of the coarse-grain model following aggregation for the system parameters plotted in a). Panels show configurations from the beginning, early, middle, and last times of the simulation, roughly times uniformly distributed along the logarithmic scale in panel a).

We have characterized the dynamics within the condensed gel state, and find that it exhibits significant dynamic heterogeneity. In Figure 5b, the mean squared displacement is plotted as a function of time for a range of temperatures within the phase separated region which shows evidence of a broad range of timescales ranging from bond breaking on the shortest timescales, to diffusive dynamics on the longest timescales. This observation aligns remarkably well with our previously reported LAT particle temporal transition from subdiffusive motion to normally diffusive motion in the LAT:Grb2:SOS condensates^[25]^. Characteristic of many glassy systems, the relaxation of a single LAT molecule is non-Markovian with significant memory of its instantaneous local gel structure. This is evidenced by the decoupling of the distribution of relaxation times we find for a LAT molecule conditioned on having just relaxed compared to its unconditioned distribution (Figure 5c). These distinct dynamical processes result in slow reorganization, which manifests long coarsening times and have origins in the limited valency, and disordered gel structure^[26]^.

## Discussion

Here we have systematically characterized kinetic aspects of the membrane-localized LAT:Grb2:SOS condensation phase transition. The results reveal that the two-dimensional phase transition and coexisting phase states differ notably from equilibrium miscibility phase separation; the LAT:Grb2:SOS phase transition is kinetically frustrated. With coarse-grained particle dynamics simulation, we identified the kinetic traps as the spindle-shaped configuration, and we observed dynamic heterogeneity in the condensed domains which are signature of glassy dynamics out of equilibrium. A recent report has identified 3D protein droplets to form soft glassy materials with age-dependent properties^[27]^. While the non-equilibrium features of this 3D system resemble some aspects of the 2D LAT system we study here, there are also some important differences as well. Most notably, although the stable steady states of the LAT system exhibit highly trapped molecular movement of LAT on the membrane surface, they are not static. The Grb2 and SOS crosslinking molecules remain quite dynamic and exchange with solution molecules through binding and unbinding^[15]^. This exchange leads to transient exposure of the underlying phosphorylated tyrosine residues on LAT, rendering them still accessible to phosphatases and kinases. Even in the condensed state, LAT molecules remain fully exposed to the solution and LAT condensates can be easily and quickly dissociated by the introduction of phosphatases at both low and high temperatures (Figure S10). In fact, the results in Figure 2c & 2d suggest that the arrested condensed phase is far from being fully-packed, which is supported by observations from the simulation (Figure 5d). This structural feature could partly be attributed to the dynamic conformational flexibility of the intrinsically disordered LAT protein.

LAT has long been known for its role as a molecular scaffold that brings together key signaling molecules once it is activated, by phosphorylation on multiple tyrosine sites. Only recently has the ensuing assembly process been recognized as a type of protein condensation phase transition, and essentially is known about what the type of phase transition may contribute to biological functionality. The results we describe here illustrate how the flexibility of LAT, along with limited valency and two-dimensionality, lead to system behavior that is predominantly kinetically controlled. In a biological setting, this suggests that the final state of a LAT condensate may be dominated by local conditions and signaling activity at the moment of its formation (for instance, kinetic competitions from other adaptor proteins^[8]^, crowdedness of the local environment, etc.) more so than the overall molecular composition and state of the cell. The low valency of crosslinking interactions between LAT molecules—with predominantly just three available sites for Grb2 interactions—plays an important role limiting LAT condensation kinetics. A fourth phosphotyrosine site on LAT, which is highly selective for PLCγ and only weakly binds Grb2, offers a fourth crosslinking possibility and has recently been reported to play an important role in stabilizing LAT condensation^[13,14]^. PLCγ is a critical signaling component in the TCR pathway, that controls Ca^2+^ signaling and ultimately T cell activation^[8,11]^. The results we report here indicate that increased valency provided by PLCγ recruitment could help break kinetic barriers limiting LAT condensation. This would ensure that LAT does not condense prior to PLCγ arrival with potentially significant consequences for coordinated signal timing (e.g. between the diverging downstream SOS and PLCγ signaling pathways). There may also be conditions under which this type of kinetic limitation is overcome in other ways, resulting in differently composed LAT condensates^[9]^. Overall, physical features of the LAT phase transition process introduce an additional angle for regulatory control and the possibility of multiple, distinct types of LAT condensate that may be able to coexist.

## Methods

### Chemicals

DOPC and DGS-NTA(Ni^2+^) were purchased from Avanti Polar Lipids. Texas red 1,2-dihexadecanoyl-*sn*-glycero-3-phosphoethanolamine (TR-DHPE) was purchased from Invitrogen. Alexa Fluor 555 maleimide dye and Alexa Fluor 488 maleimide dye were purchased from Life Technology. BSA, (±)-6-hydroxy-2,5,7,8-tetramethylchromane-2-carboxylic acid (Trolox), catalase, 2-mercaptoethanol (BME), NiCl2, H2SO4, and ATP were purchased from Sigma-Aldrich. Glucose oxidase was purchased from Serva. Tris(2-carboxyethyl)phosphine (TCEP) was purchased from Thermo Scientific. Glucose and H2O2 were from Fisher Scientific. MgCl2 was from EMD Chemicals. Tris buffer saline (TBS) was purchased from Corning.

### Protein Purification and Labeling

LAT, Grb2 and SOS_PR were purified similarly to previously published works^[7,15,25]^. Human LAT cytosolic domain (residues 27 to 233, C117S) and full-length Grb2 were purified as described previously using an N-terminal 6-His tag, while proline rich domain of SOS (residue 1051-1333) was purified with similar strategy. For both Grb2 and SOS_PR, the N-terminal 6-His tag were removed with Tobacco Etch Virus protease.

Protein was diluted to 50 μM, and Alexa Fluor dyes were prepared at 10 – 20 mM in anhydrous dimethyl sulfoxide (DMSO). The protein was incubated with 5 mM TCEP on ice for 30 minutes to ensure all cysteines were reduced. The dye was then added in 1 – 5 fold molar excess, depending on the percentage of labeling desired and the stability of the protein in the presence of the free dye. The proteins were then allowed to react with the dye for 2 h on ice. The reaction was then quenched with 10 mM DTT for 10 min. Excess dye was removed by Superdex 75 column (10/300 GL, GE Healthcare). Percent labeling was calculated by measuring absorbance at peak excitation for the dye and 280 nm for the protein, while taking into account 280 nm contribution from the dye. Labeled protein was concentrated to the desired concentration, flash frozen, and stored at -80°C.

### Formation of Functionalized Supported Lipid Bilayers

Glass substrates (no. 1.5 thickness) were prepared by 10 min sonication in 1:1 IPA/water mixture, 90s microwave heating in water, 30 min soaking in 2 v% Helamax solution and 5 min of piranha etching (H2SO4:H2O2 = 3:1 by volume), followed by excessive rinsing of H2O (Milli-Q). The thus cleaned glass substrates were stored in Milli-Q water for no more than three days before using. The substrates were blown dried with air before depositing vesicles to form supported lipid bilayers (SLBs).

Small unilamellar vesicles (SUVs) were prepared by mixing DOPC:Ni^2+^-NTA-DOGS = 96:4 by molar percent in chloroform. If visualization of the bilayers was required, additional 0.005 mol% of TR-DHPE was added into the lipid composition. The solution mixture was then evaporated by a rotary evaporator for 15 min at 40 °C. Lipid dried films were further blown with N2 for another 15 min. The lipids were then resuspended in H2O by vortexing, resulting in a concentration of about 1 mg/mL. Lastly, the vesicle solution was sonicated with a tip-sonicator for 5 min with 20 s-on and 50 s-off cycles in an ice-water bath. The membrane system was prepared on a flow chamber (μ-Slide; Ibidi) for experiments at and below 25 °C. For experiments at temperatures higher than 25 °C, the SLBs were formed in a PDMS ring attached to a round glass coverslip, which then brought into an incubator and incubated at the desired temperature. SLBs were formed on the glass substrate by incubating the SUVs mixed 1: 1 v/v with 2X TBS (10 mM Tris and 150 mM NaCl, pH 7.4) for at least 30 min. The chambers were rinsed with 1X TBS buffer and then incubated with 100 mM NiCl2 for 5 min. Next, 1 mg/mL BSA was incubated for 10 min to block defects in supported membranes. Before protein incubation, the system was buffer-exchanged into protein dilution (PD) buffer containing 1X TBS, 5 mM MgCl2 and 1 mM TCEP, pH 7.4. Proteins were centrifuged for 20 min at 4 °C beforehand to remove possible aggregates. Depending on the desired density of the membrane proteins, LAT (50 nM – 500 nM) and Hck (15 nM) were incubated for 45 min to attach onto the bilayers via His-tag chemistry^[28]^. The system was allowed to sit for another 20 min for unstably bound membrane proteins to dissociate from the surface. Between all incubation steps, the chambers were rinsed with PD buffer. All parts of preparation were done at room temperature unless stated otherwise.

### Microscopy

TIRF experiments were performed on a motorized inverted microscope (Nikon Eclipse Ti-E; Technical Instruments, Burlingame, CA) equipped with a motorized Epi/TIRF illuminator, motorized Intensilight mercury lamp (Nikon C-HGFIE), Perfect Focus system, and a motorized stage (Applied Scientific Instrumentation MS-2000, Eugene, OR). A laser launch with 488 and 561 nm (Coherent OBIS, Santa Clara, CA) diode lasers was controlled by an OBIS Scientific Remote (Coherent Inc., Santa Clara, CA) and aligned into a fiber launch custom built by Solamere Technology Group, Inc. (Salt Lake City, UT). The optical path was then aligned to a 100x 1.49 NA oil immersion TIRF objective (Nikon). A dichroic beamsplitter (ZT405/488/561/640rpc; Chroma Technology Corp., Bellows Falls, VT) reflected the laser light through the objective lens and fluorescence images were recorded using an EM-CCD (iXon 897DU; Andor Inc., South Windsor, CT) after passing through a laser-blocking filter (ZET405/488/561/640m-TRF; Chroma Technology Corp., Bellows Falls, VT). Laser powers measured at the sample were approximately 8 mW for the 488 nm channel when taking the single molecule tracking measurements. All acquisitions were obtained using Micro-Manager. A TTL signal from the appropriate laser triggered the camera exposure.

### Single Particle Tracking

All analysis was performed on the central region of the images with radius of 160 pixel to minimize uneven illumination. Particle identification and particle tracking routines were performed using Trackmate. Single particle was identified and detected by a segmenter based on an approximation of the Laplacian of Gaussian (LoG) operator by difference of gaussian (DoG), with sub-pixel localization and an estimated particle diameter of 0.6 μm. An initial quality thresholding was performed before tracking analysis to ensure optimized single particle density, which in this case is 0.07 molecules/μm^2^ to 0.1 molecules/μm^2^. Particle diffusion was tracked by the Linear Assignment Problem (LAP) tracker, with the maximum travel distance of 16 pixel per frame at framerate of 33 Hz and maximum frame gap of 2 frames. All results were inspected to verify proper particle connection, and only tracks displaying more than five spots are processed for further analysis.

### Simulation Methods

Each LAT molecule is treated as a disk with a diameter of σ = 5 nm based on the radius of gyration. The dynamics of the *i*’th LAT follows an overdamped Langevin equation of the form,

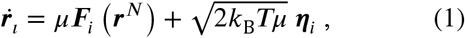

where is *k*_B_ Boltzmann’s constant, μ is the single particle mobility, *T* is the temperature of the system, and combined they set the temperature dependent diffusion by the fluctuation-dissipation relation in the dilute limit, *D* = *k*_B_*T*μ. The interactions, *F*_*i*_(*r*^*N*^) = −∇_*i*_*U*(*r*^*N*^), are conservative, pairwise additive, and in general depend on the coordinates of all *N* particles in the system represented by configuration *r*^N^. The statistics of *η*_*i*_ are zero on average, 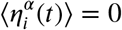, and have a variance that is delta correlated in time, 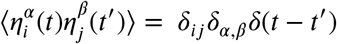, for the *α* and *β* components of the random force which correspond to the *x* and *y* dimensions. In practice, the mobility and Boltzmann’s constant are set to unity and the dynamics are generated a first order Euler method with Ito time discretization. We consider conservative pairwise forces with excluded area given by the Weeks-Chandler-Anderson potential^[29]^,

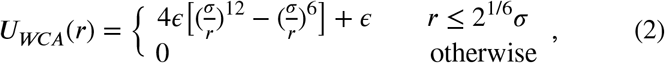

with a characteristic energy scale *ϵ* = 1. Bonded LAT molecules also have a pairwise bond force with other LAT molecules given by a harmonic potential,

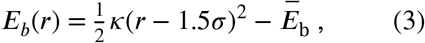

where Ē_b_ is set to 4*ϵ*. A given LAT can at most be bound to three other LAT molecules but the number of bonds at a given time is a dynamic quantity since we allow for bonds to be made and broken. Bonds between pairs of LAT are formed and broken with Metropolis-Hastings algorithm with an energy change to make a bond given by *E*_*b*_(*r*) and to break a bond given by −*E*_*b*_(*r*) [30, 31]. A bond is made at rate *k*_*b,r*_ when two LAT molecules are within a reactive distance of 2σ and have less than three bonds. A bond is broken with rate *k*_σ,*r*_ given that the pair of LAT molecules are within a reactive distance of *r*_*c*_ = 2σ. The rates are related to each other by a detailed balance relation,

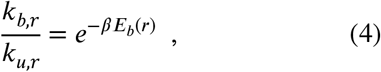

with

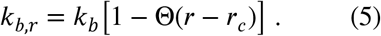

Here *k*_*b*_ is the prefactor rate which sets the time scale of the kinetics and Θ(*r* − *r*_*c*_) is a Heaviside function that enforces that only when particles are within a reactive distance are they capable of forming a bond. To ensure realistic dynamics, we choose a tight spring constant of *Κ* = 80*ϵ*/ σ^2^ to avoid bound LAT extending beyond the reactive distance.

The simulation code used can be found here: https://github.com/tgrandpr/LAT_simulations.

## Acknowledgement

This work was supported by the Novo Nordisk Foundation Challenge Program under the Center for Geometrically Engineered Cellular Systems. Additional support was provided by National Institutes of Health grant PO1 A1091580. T. G. P and D. T. L were supported by the National Science Foundation, Division of Chemistry Award No. NSF Grant CHE1954580. We thank L. B. Nocka, L. J. N. Lew, A. A. Lee and J. B. DeGrandchamp for sharing reagents and helpful discussions.

## Author contributions

S. S. and J. T. G. conceived the study. S. S. designed and performed the experiments, analyzed the experimental data. T. G. and D. T. L. developed a model, ran simulations, and analyzed the data from it. S. S., T. G., D. T. L. and J. T. G. wrote the manuscript.

