## Supplemental materials for "Kinetic frustration by limited bond availability controls the LAT protein condensation phase transition on membranes"

LAT density calibration and mapping the phase diagram:

All TIRF images were processed by flattening and background subtraction before analysis. For a homogeneous LAT layer, a calibration curve between the surface density of LAT measured by FCS and the corresponding TIRF intensity was first made (Figure S3). Based on the linear correlation, the pixel brightness histogram of LAT distribution can then be calibrated to LAT density distribution as shown in Figure 1.

To map out a phase diagram for the LAT:Grb2:SOS phase transition as demonstrated in Figure 2c, after the phase transition reached the apparent final state (no detectable change in the normalized intensity variance), the LAT density distribution was fitted with two Gaussian peaks. The high-density peak position represents the average LAT density in the condensed phase, and the low-density peak position represents the average LAT density in the dispersed phase. The same strategy was carried out at varied temperatures, and the density difference normalized to the initial protein density (1200 ± 600 molecules/μm$^2$) was mapped out.

To validate the above analysis, a different scheme was taken to map the apparent final state phase diagram. First, we fitted the pixel brightness histogram with two Gaussians. Next, we used the high-intensity peak position ($F_c$) and the low-intensity peak position ($F_d$) as the average intensity of the condensed phase and the dispersed phase respectively. In this way, the respective area fractions of the condensed phase ($A_c$) and the dispersed phase ($A_d$) can be calculated with intensity-based image segmentation. Lastly, we applied the following set of equations to calculate the average LAT density in the two phases ($x_{c,avg}$ and $x_{d,avg}$):

$$x_{c,avg} \times A_c + x_{d,avg} \times A_d = x_0 \times A_{total} \quad (Eqn.\ S1)$$

$$A_c + A_d = A_{total} \quad (Eqn.\ S2)$$

$$\frac{x_{c,avg}}{x_{d,avg}} = \frac{F_c}{F_d} \quad (Eqn.\ S3)$$

The resulting phase diagram across different temperatures (Figure S4a) is similar to Figure 2c.

It is worth noting that, when conducting the intensity-based image segmentation according to the two-Gaussian fitting of the pixel brightness histogram, in a couple of trials, we observed that some scattered, small and less dense condensed domains were segmented as the dispersed phase. To examine how much this effect might influence the apparent final state phase diagram, we performed three-Gaussian fitting to the LAT density distribution profile. By fixing the middle peak position to be the initial (average) LAT density, the less dense small clusters are now incorporated into the middle peak and are separated from both the condensed phase peak and the dispersed phase peak. A phase diagram obtained in this way (Figure S4b) is also similar to Figure 2c.

Analysis of the phase transition kinetics:

Due to the linear relationship between LAT protein surface density and TIRF intensity, we used the normalized variance of the fluorescence intensity to characterize the normalized variance of the LAT density distribution during the phase transition. The normalization was made to the average intensity of the images, and the normalized variance represents the normalized mean squared LAT density fluctuations ($\langle (dx)^2 \rangle$). By tracking the normalized mean squared LAT density fluctuation, we are able to extract the kinetic profile of a LAT:Grb2:SOS phase transition.

The kinetic profile was fitted to a sigmoidal function as follows:

$$y = \langle (dx)^2 \rangle_{max} + \frac{\langle (dx)^2 \rangle_{lag} - \langle (dx)^2 \rangle_{max}}{1 + e^{(t-t_{50})\times \frac{4k}{\langle (dx)^2 \rangle_{lag} - \langle (dx)^2 \rangle_{max}}}} \qquad (Eqn.\, S4)$$

where the slope $k$ at $t = t_{50}$ was extracted to represent the rate of the growth stage of the phase transition, and $t_{lag}$ presented the length of the time delay between adding linker proteins and the start of the growth stage. $\langle (dx)^2 \rangle_{max}$ was consistently set to 1, and in the majority of the traces, $\langle (dx)^2 \rangle_{min}$ was at zero. However, in about 1 out of 5 trials, we observed $\langle (dx)^2 \rangle_{min}$ bigger than zero, suggesting that there was detectable LAT density fluctuation before the start of the growth stage. In these cases, the small increase in the normalized variance during the lag stage was well fitted to the Avrami model in Eqn. S5[1]:

$$y = \langle (dx)^2 \rangle_{lag} \times \left(1 - e^{(-(mt)^n)}\right) \qquad (Eqn.\, S5)$$

where *m* is a constant and *n* is the so-called Avrami or stretch exponent. The good fit to the Avrami model indicates that the stochastic and dynamic small density fluctuation in the lag stage was likely due to heterogeneous nucleation events.

Further details on the simulation methods:

All simulations are done with a system size of $N = 1000$, a time step of $\delta t = 10^{-4}$, and observation times ranging from $10^8 \delta t - 10^9 \delta t$. The phase diagram as a function of density and temperature is shown in Figure S7a. The critical temperature depends on the minimum bonding energy, $\overline{E}_{\rm b}$, which is set to $4\epsilon$ for all simulations. At that minimum bonding energy, the critical temperature is at about $k_B T / \overline{E}_{\rm b} \approx 0.19$ and a reduced density of $\rho\sigma^2 \approx 0.16$. A system with parameters in the orange shaded region will be phase separated into a condensed and dispersed phase with average density represented by the light and dark blue squares with error bars obtained from the standard error from two different methods. Outside of that region, the system will be homogeneous. One method to obtain the coexisting densities was to fit a slab embedded in a rectangular system of dimensions $L_x = 6L_y$ to a sigmoidal function. The other method used was a subsystem analysis where a probe area was moved around the system and a bimodal distribution peaked at the coexisting densities was gotten, $P(\rho_a)$, for probe area $a = \pi d^2$. In both methods, a system density of $\rho\sigma^2 = 0.1$ was used. The subsystem density distributions above, below, and around the critical point are shown in Figure S7b) at a total system density of $\rho\sigma^2 = 0.1$. Below the critical point, the distribution is bimodal at the coexisting temperatures in agreement with Figure S7a. Near the critical point, the difference between the peaks shrinks and the bimodality is lost. Above the critical point, the distribution becomes narrow but is still non-Gaussian. In Figure S7c), for $k_B T / \overline{E}_b \approx 0.18$ the average density, $\langle \rho_a \rangle_{\Delta\mu}$, is plotted as a function of the relative chemical potential between the condensed and dispersed phases, Δμ, computed by reweighting the density distribution as

$$\langle \rho_a \rangle_{\Delta\mu} = \frac{\langle \rho_a e^{a\beta\Delta\mu\rho_a} \rangle_0}{\langle e^{a\beta\Delta\mu\rho_a} \rangle_0} \quad , \qquad (Eqn.\, S6)$$

where the average is taken over the subsystem distribution, $P(\rho_a)$. The critical chemical potential is a value close to zero but positive. For a relative chemical potential above the critical value, the density increases and then approaches the condensed phase at $\rho\sigma^2 \approx 0.38$. For a relative chemical potential below the critical value, the density decreases and eventually reaches the dispersed phase density at $\rho\sigma^2 \approx 0$. The intensive variance exhibits finite size effects with the peak growing larger with increasing probe area size. The intensive variance can be obtained from by taking a derivative of $\langle \rho_a \rangle_{\Delta\mu}$with respect to Δμ,

$$a\langle (\Delta\rho)^2 \rangle_{\Delta\mu} = \frac{1}{\beta}\frac{d\langle \rho_a \rangle \Delta\mu}{d\Delta\mu} \quad . \qquad (Eqn. S7)$$

which is shown in In Figure S7d.

Within our work, we also study the steady state and dynamical properties of the isolated dense phases represented by the blue squares in Figure S7a. The dynamical properties are shown in the main text but here we describe the steady state properties in more detail in Figure S9. In Figure S9a, the radial distribution function,

$$g(r) = \frac{1}{\rho N}\langle \sum_{i \neq j}^{N} \delta(r - r_{ij}) \rangle \quad , \qquad (Eqn. S8)$$

is plotted as a function of the average distance between particles, $r_{ij} = \sqrt{(x_i - x_j)^2 + (y_i - y_j)^2}$. It is seen that the distribution function looks liquid-like with some short ranged correlations but no long range order. In Figure S9b, a normalized histogram of the relative angle between two bonded LAT molecules labelled 1 and 2 of the $i$'th LAT molecule,

$$\theta = cos^{-1}\left(\frac{\boldsymbol{r}_{i1} \cdot \boldsymbol{r}_{i2}}{r_{i1} r_{i2}}\right) , \qquad (Eqn. S9)$$

is plotted for the temperatures of $k_B T / \bar{E}_b \approx$ 0.18, 0.15, and 0.14 and their corresponding dense phases of $\rho\sigma^2 = 0.27, 0.32$ and 0.36. When a given LAT has three bonds there are three different angles that enter the histogram and there are no angles recorded when LAT has less than two bonds. For the temperatures studied, there is a peak value at about 1.2 radians which gives rise to the spindle-like structure shown in Figure 5 of the main text.

We use the knowledge of the phase diagram to understand growth and the dynamics of the dense phase. In Figure 5a of the main text, a system at $k_B T / \bar{E}_b = 0.5$ and $\rho\sigma^2 = 0.1$ was quenched to $k_B T / \bar{E}_b = 0.13$ at the same system density. The largest cluster at each time slice was determined using a cluster algorithm. On the right axis of the same plot, the average bonds is calculated by dividing the total bonds by $N$ at each time slice. This is also averaged over 11 runs with negligible error bars.

For the temperatures $k_B T / \bar{E}_b \approx$ 0.18, 0.15, and 0.14, we ran simulations with densities corresponding to dense phases coexistence densities, $\rho\sigma^2 = 0.27, 0.32$ and 0.36, and measured the mean squared displacement,

$$\langle \Delta r^2(t) \rangle = \frac{1}{2N}\sum_{i=1}^{N} |r_i(t) - r_i(0)|^2 \ . \qquad (Eqn. S10)$$

These are illustrated in Figure 5b of the main text.

Estimation of bond energy between each pair of LAT molecules:

We consider the bond formation between each pair of LAT molecules by the interactions between pLAT (fully phosphorylated LAT) and Grb2, Grb2 and SOS_PR, then SOS_PR to another Grb2, which eventually binds to the second pLAT. The binding affinity between a fully phosphorylated LAT and the SH2 domain of Grb2 was estimated to be 10 nM – 100 nM [2]. The interaction between Grb2 and SOS_PR can be more complicated, considering that there are at least four binding sites on SOS_PR that can bind to the SH3 domains of Grb2, and the full length Grb2 protein can bind either monovalently or bivalently to SOS_PR. As such, the binding affinity of Grb2 and SOS_PR has a wide range of 10 nM to 1 mM[3,4]. Therefore, the calculated bond energy between each pair of LAT molecules is in the range of 46 $k_{\mathrm{B}}T$ – 73 $k_{\mathrm{B}}T$ (at 298 $K$).

Supplementary figures:

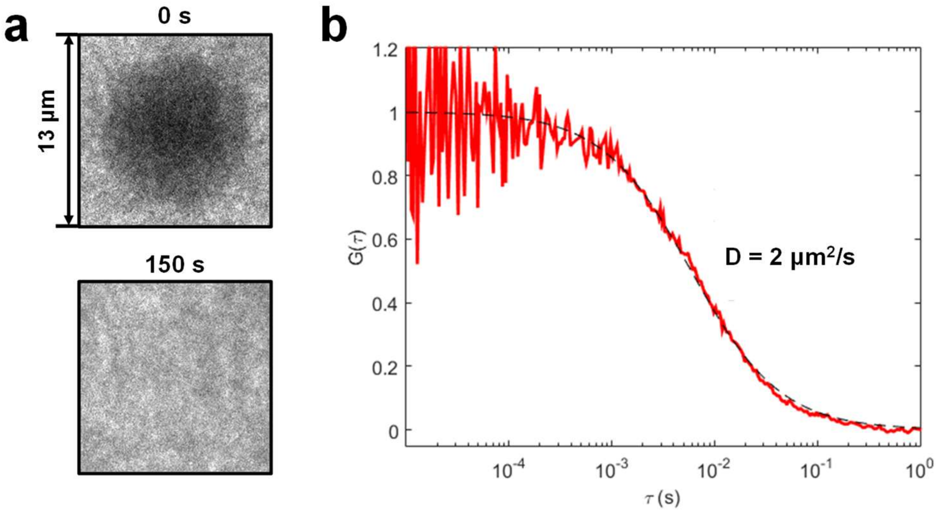


Figure S1. Characterization of the membrane-bound LAT proteins before phase transition: a) FRAP results of the homogeneously distributed LAT layer, with the upper image taken right after photobleaching and the lower image taken 150 s later which demonstrates close to full fluorescence recovery. b) Representative FCS trace of monomeric diffusing LAT (red solid curve) fitted to a two-dimensional Gaussian diffusion model (black dashed line), with a fitted diffusion coefficient of D = 2 $\mu m^2/s$.

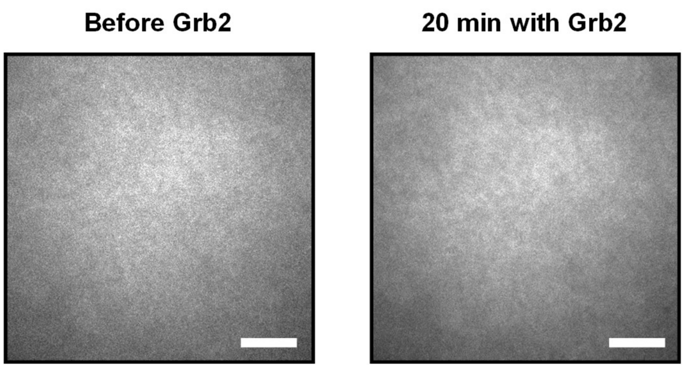


Figure S2. TIRF images of the LAT layer before and after 20 minutes incubation with 5.8 μM Grb2. Scale bar: 7 μm.

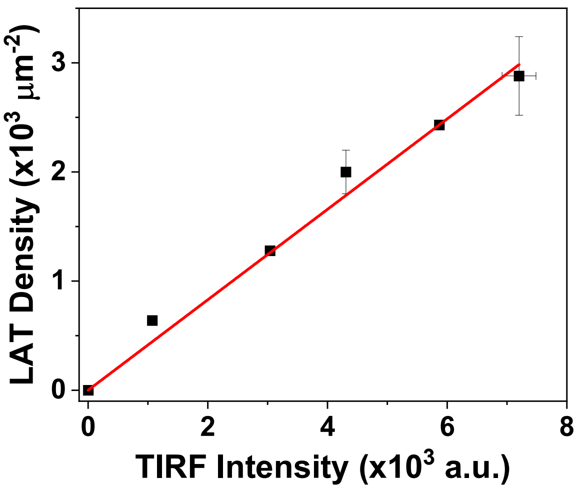


Figure S3. Calibration curve between TIRF intensity of the LAT layer and the LAT surface density measured by FCS. The SD is calculated based on three individual measurements.

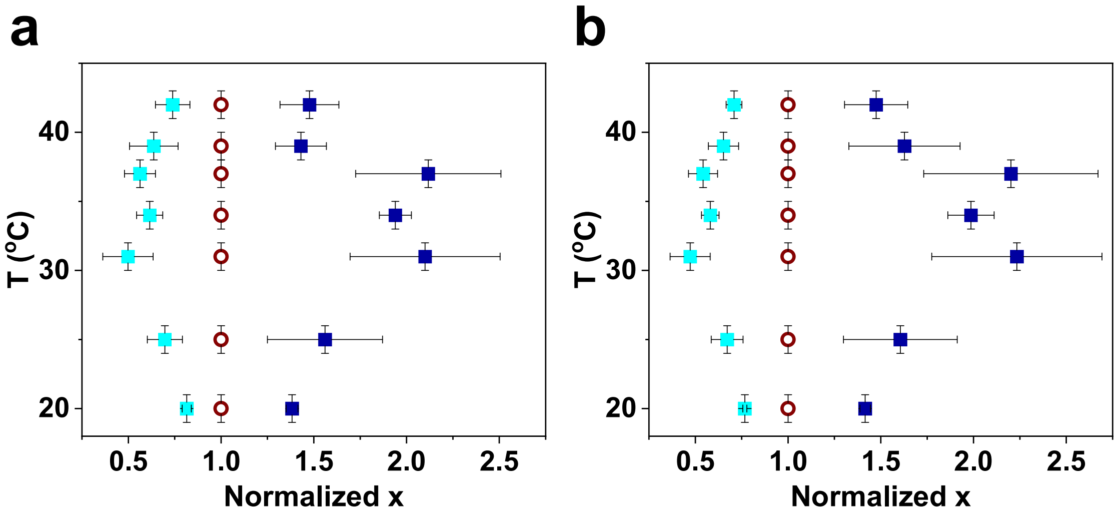


Figure S4. Phase diagrams at the apparent final state plotted based on two different analysis schemes: a) Pixel brightness histogram was first fitted to two Gaussian peaks, then corresponding LAT density in the two phases was calculated according to the segmented area fractions. b) The LAT density distribution was fitted to three Gaussian peaks. Red open circle: initial LAT density before phase transition, royal blue solid square: average LAT density in the condensed phase, cyan solid square: average LAT density in the dispersed phase.

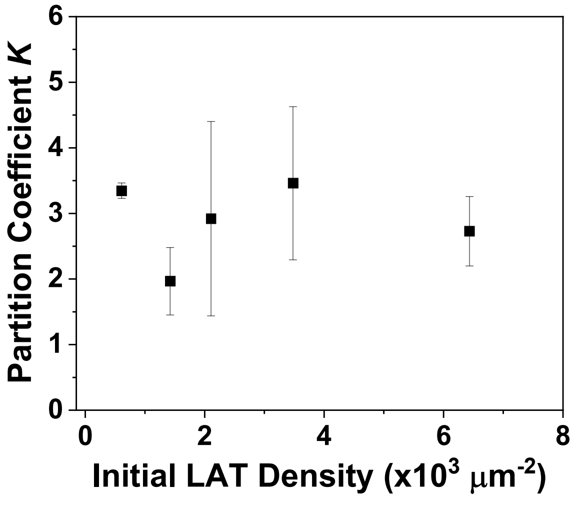


Figure S5. Partition coefficient $K$ as a function of initial LAT density.

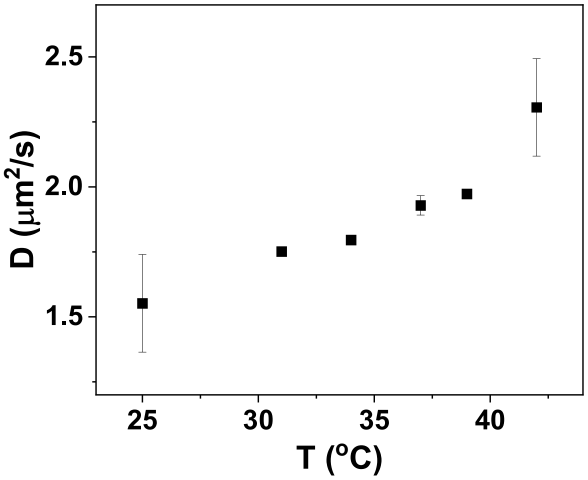


Figure S6. Diffusion constant of the membrane-anchored LAT monomers in the temperature range from 25 °C to 42 °C measured by both FCS and FRAP.

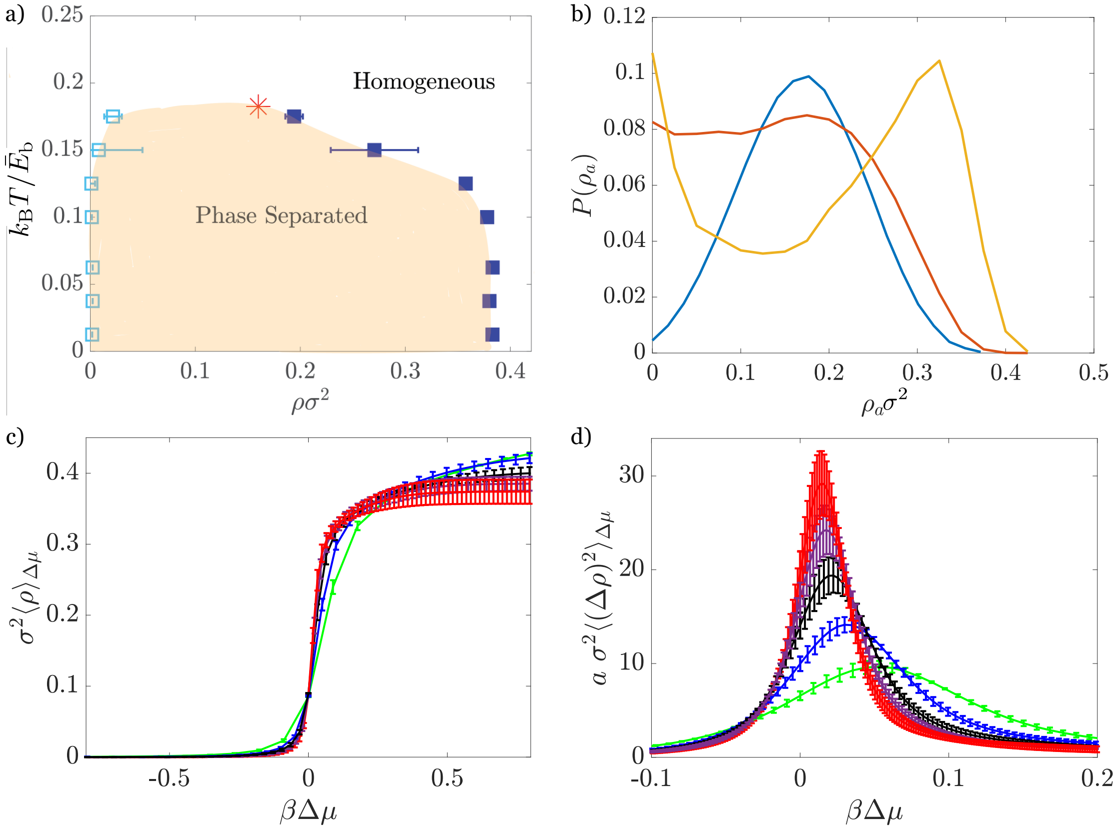

Figure S7: Phase diagram of coarse-grained LAT model. a) The phase diagram as a function of the reduced density and reduced temperature. The light blue squares are the dilute phase points and the solid blue squares are the dense phase points. The shaded region represents where there is phase separation. b) The distribution of subsystem densities, reweighted by the critical chemical potential density denoted by the peak in the variance, for temperatures below, around, and above the critical temperature at $k_B T/\bar{E}_b = 0.13$ (yellow), 0.18 (red), and 0.38 (blue) with a probe radius of $8\sigma$ and $\rho\sigma^2 = 0.1$. c) The average density from $Eqn. S6$ as a function of the relative chemical potential between the dense and dilute phase, Δμ, for circular probe areas of radii $d = 6\sigma$ (cyan), $8\sigma$ (blue), $10\sigma$ (black), $12\sigma$ (purple), and $14\sigma$ (red) at $k_B T/\bar{E}_b = 0.18$ and $\rho\sigma^2 = 0.1$. d) The average variance from $Eqn. S7$ as a function of the relative chemical potential for the same subsystem sizes as b).

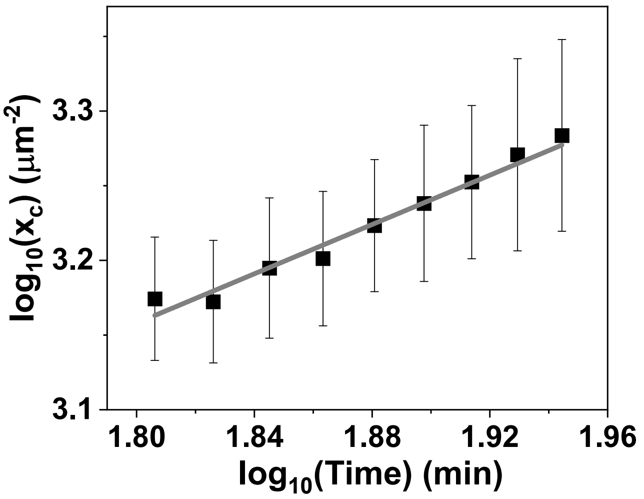


Figure S8. Average LAT density in the condensed phase ($x_c$) as a function of time plotted in the log-log scale. The fitted slope of the linear trendline is 0.8.

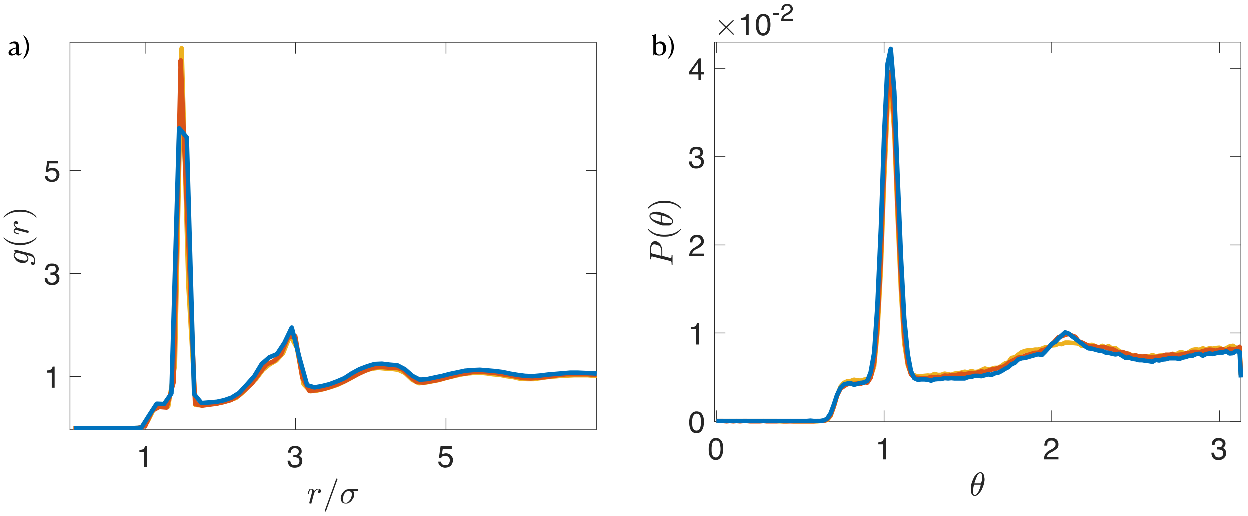

Figure S9: Steady state properties. a) The radial distribution function for temperatures of $T/\bar{E}_b = 0.18$ (blue), 0.15 (red), and 0.14 (yellow) and their corresponding dense phase densities of $\rho\sigma^2 = 0.27$, 0.32, and 0.36. b) The distribution of relative angles between two bonds of a LAT molecule with the same parameters as b) with an average at 1.2 radians (68 degrees).

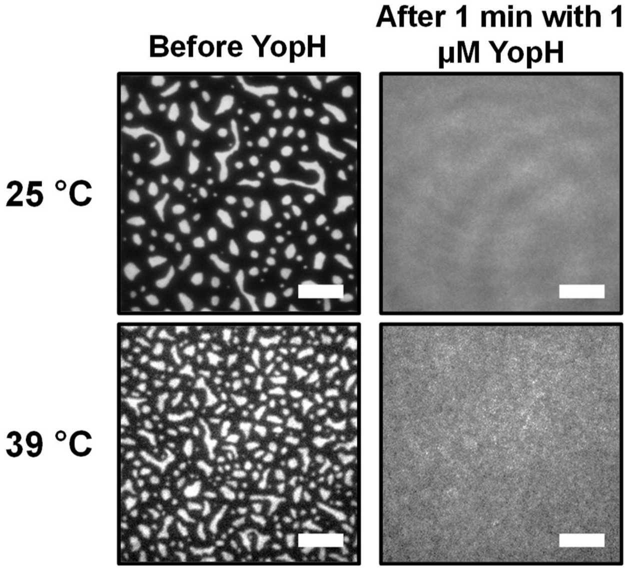


Figure S10. Quick disassembly of the LAT:Grb2:SOS condensates by a promiscuous phosphatase YopH at 25 °C and 39 °C. Scale bar: 7 μm.

Movie S1. Demonstration of the dynamic fluctuations at the edges of the condensed domains. The movie was taken at a time interval of 20 s, after the phase transition reached the apparent final state. Scale bar: 2 μm.
